# Identification and functional characterization of *LDB3* gene variants in DCM patients from Indian Population

**DOI:** 10.1101/2025.10.21.683814

**Authors:** Bhagyalaxmi Mohapatra, Amrita Mukhopadhyay, Prerna Giri, Dharmendra Jain, Ashok Kumar

## Abstract

The LIM domain-binding protein 3 (LDB3), also known as Z-band alternatively spliced protein (ZASP), is a key component of the Z-disc, essential for maintaining sarcomeric integrity and transmitting contractile force in cardiomyocytes. Genetic variants in LDB3 have been implicated in cardiomyopathies, including dilated cardiomyopathy (DCM). In this study, we have screened 100 idiopathic DCM cases along with controls by Sanger sequencing and identified nine non-synonymous (p.S184I, p.D193N, p.K204R, p.R229C, p.P295T, p.Q402P, p.F465I, p.F496L, p.P606S), seven synonymous (p.T91=, p.A152=, p.D168=, p.S182=, p.A279=, p.S347=, p.A358=), and seven intronic (c.322-50G>C, c.548+53A>C, c.718+47G>C, c.718+81G>A, c.755+11G>A, c.755+86G>A, c.*30C>G) variants, among which six (p.S184I, p.D193N, p.P295T, p.F465I, p.F496L, p.P606S) missense variants were novel. *In-vitro* studies demonstrated that p.K204R increased LDB3 expression whereas p.S184I, p.F465I, p.F496L, p.P606S reduced LDB3 expression. Immunostainning revealed cytoplasmic aggregation of the p.K204R-LDB3 protein. The variant p.R229C also over-expressed the mutant protein. In case of variant, p.P295T, Z-disc was severely disrupted indicating impaired Z-disc integrity. Furthermore, synonymous variants showed altered mRNA folding, stability, and codon usage bias, potentially affecting translation efficiency. In-silico analyses predicted p.F465I, p.F496L, p.P606S, and p.S184I variants to be deleterious, significantly altering protein structure. Structural modeling using AlphaFold2 showed high RMSD values, suggesting conformational destabilization. Collectively, these findings highlight LDB3 as a hypermutable candidate gene for DCM in our cohort that may contribute to DCM pathogenesis by perturbing protein structure, expression, and cellular localization, underscoring the critical role of LDB3 in cardiac muscle function and disease.

## 1 Introduction

Z-disc is a multi-protein structure, seen as electron-dense band with electron microscopy, harbor a plethora of proteins, forming lateral boundary of sarcomere. These proteins play a pivotal role in signal transduction within cardiomyocytes and are also important for cross-linking several proteins along with transmission of force generated by the myofilaments for contraction of cardiac muscles (Frank *et al*. 2006). One of these proteins, that has been shown to provide tensile strength to the cardiomyocytes, is the LIM domain binding3 protein (LDB3), also known as Z-band alternatively spliced protein (ZASP).

LBD3 is a member of Enigma family of proteins, predominantly expressed in cardiac and skeletal muscles (Lopez-Ayala *et al*. 2015). The mouse homologue of LDB3, Cypher has been cloned and identified as a striated muscle restricted PDZ/LIM domain protein which colocalizes with α-Actinin-2 (ACTN2) at the Z-disc (Zhou *et al*. 1999). LDB3 has been independently identified in heart and skeletal muscle (Faulkner *et al*. 1999). Due to alternative splicing LDB3 has eight isoforms in humans. The transcript variants 1,2,5,6 and 8 express in adult heart. All these isoforms also express in adult skeletal muscles.

The protein encoded by longer transcript variants (transcript variant 1 and 2) of LDB3 in humans have PDZ domain (5-81 amino acids), domain of unknown function (DUF4749; 148-238 amino acids), Atrophin-1 domain (272-439 amino acids) and three LIM domains (LIM1; 441-492, LIM2; 500-551, LIM3; 551-612) towards C-terminus. Additionally, in transcript variant 5, a ZM (ZASP like) motif is present which interacts with the spectrin repeats of ACTN2 (Klaavuniemi *et al*. 2004; Klaavuniemi and Ylanne 2006). The PDZ domain of LDB3 is present in isoforms 1, 2, 3, 4, 5, 6, 7 whereas LIM domain is present only in isoforms 1, 2 and 5. The LIM domains of LDB3 interact with protein kinases (Pathak *et al*. 2021) and mediate diverse protein-protein interactions, including the interaction with Calsarcins. Studies using human embryonic kidney-293 cells and neonatal rat cardiomyocytes, have shown that the PDZ domain of LDB3/ZASP binds to Telethonin (T-CAP) forming a complex with the sodium channel Na(v)1.5 (Xi *et al*. 2007). LDB3 also binds to Nebulin (NEB) and Phosphoglucomutase 1 (PGM1). Molecular genetic studies have indicated that variants in *LDB3* have been associated with myopathies including different forms of myopathies such as DCM, HCM and LVNC (Vatta *et al*. 2003; Arimura *et al*. 2004). Mutations in this gene have also been associated with myofibrillar myopathy (Griggs, *et al*. 2007; Lin *et al*. 2014).

The first ever evidence for the role of *LDB3* in cardiomyopathy came from the studies on *Cypher* knock-out mouse. The knock out animals developed severe forms of congenital myopathy and died early, most likely due to cumulative effect of cardiomyopathy and respiratory failure. *Cypher* is required for maintaining Z-disc integrity and it has been proposed to be a ‘linker-strut’ in contracting muscles (Zhou *et al*. 2001). Deletion of *cypher* resulted in premature death and severe DCM in both post-natal and in adult heart, with altered cardiac signaling and disorganization of Z-disc (Zheng *et al*. 2009). A zebrafish morphilino knockdown of *cypher* also resulted in DCM (Meer *et al*. 2006). In humans, *ZASP* associated mutations were reported in DCM and LVNC cases for the first time by Vatta *et al*. 2003. The mutation (D117N) identified in this studied indicated cellular disarray as revealed by in vitro transfection assays. In another study exploring the mechanism by which a previously published S196L mutation leads to the phenotype of DCM, it was found that in the cardiomyocytes isolated from S196L transgenic mice, the calcium and sodium channels were altered (Li *et al*. 2010). Parallely, Arimura *et al*. 2004 also identified DCM-associated mutation (D626N) in *ZASP* in Japanese population. This mutation increased the binding affinity of ZASP with PKC as evident by *in-vitro* protein-protein interaction studies. A very recent study using *in vitro* approaches, reported that *Cypher* deficiency induces apoptosis (Xuan *et al*. 2020) and this finding was consistent with the fact that Cypher acts as a negative regulator of the proapoptotic protein TP53 (Martinelli *et al*. 2014).

Accumulating evidences using animal models and human genetic studies have suggested a pivotal role of LDB3 for maintenance of cardiac structure and function. However, the mechanism underlying the *LDB3* mediated DCM phenotype is not clearly understood. Further, molecular genetic investigations are required to provide precise insights into association of DCM with *LDB3*. This prompted us to perform a detailed molecular genetic analysis of *LDB3* in a cohort of 100 DCM cases. We identified a handful of variants in *LDB3* which included nine non-synonymous, seven synonymous and seven intronic variants that were not found in hundred controls from our population suggesting the possible disease-association of these variants. We observed a high mutation frequency of 21% in the protein-coding region. The non-synonymous variants were characterized using *in vitro* approaches indicating that *LDB3* is required for maintaining the integrity of Z-disc and we also characterized synonymous variants using computational approaches which indicated altered mRNA conformation and stability and also codon bias.

## 2 Materials & Methods

### 2.1 Clinical Evaluation and Enrolment of Patients

A total of one hundred clinically diagnosed idiopathic DCM patients (median age at diagnosis of 11 years) were recruited from the Department of Pediatric Medicine and Department of Cardiology, Institute of Medical Sciences, Banaras Hindu University, along with 100 age and ethnic-matched healthy individuals as controls (median age 3.7 years) for the present study. The diagnosis of patients was based on physical examination, chest radiography, MRI, ECG and 2D-echocardigraphy for left ventricle (LV) size and function (i.e., LV fractional shortening <25%, an ejection fraction of <45%). The study was approved by the ‘Institutional Ethics Committee’ for involving humans as research subjects. A prior written informed consent was obtained from all the patients or their authorized guardians in order to collect blood samples for genetic investigations.

### 2.2 Mutational Analysis of *LDB3*

#### 2.2.1 Genomic DNA isolation

For genetic investigations, peripheral venous blood samples (3-5 ml) were collected from the patients, their available relatives, and unrelated healthy control individuals, after written informed consent. Genomic DNA was extracted according to the standard ethanol precipitation protocol. The completely dissolved DNA was checked qualitatively and stored at 4°C till further use.

#### 2.2.2 Polymerase Chain Reaction

All 16 protein-coding exons and exon-intron boundaries of the longest isoforms of human *LDB3* (NM_001080114.2; isoform 2 and NM_007078.3; isoform 1, together compose all 16 exons) were amplified by polymerase chain reaction (PCR) using sets of forward and reverse primers designed at respective exon-intron boundaries. All amplified products were purified using 3U of Exonuclease I (USB Products, Affymetrix, Inc, OH, USA) and 0.6 U of Shrimp Alkaline Phosphatase (USB Products, Affymetrix, Inc, OH, USA) by incubating at 37°C for 45 minutes, following which enzymes were heat-inactivated at 80°C for 15 mins.

#### 2.2.3 Sequencing of PCR products by Sanger’s Method

The purified PCR products were subjected to Sanger’s sequencing using Big Dye Terminator sequencing kit 3.1 (ABI, USA) as per the manufacturer’s protocol. Sequencing was performed on 3130 Genetic Analyzer, (Applied Biosystems). Finch TV software (http://www.geospiza.com/ftvdlinfo.html, Geospiza) was used to analyze the chromatograms. Each identified variant was confirmed by re-sequencing of freshly amplified PCR products of the same amplicon from the DNA of respective subjects. Novelty of the sequence variants was checked by Genome Aggregation Database (gnomAD) (https://gnomad.broadinstitute.org/) and regional databases such as INDEX-db (http://indexdb.ncbs.res.in/), IndiGenomes-DB (https://clingen.igib.res.in/indigen/) and GenomeAsia100K (https://browser.genomeasia100k.org/).

### 2.3 *In silico* characterization of identified variants

#### 2.3.1 Phylogenetic conservation of amino acid residues in LDB3 protein

The reference genomic DNA sequence, mRNA sequence and protein sequence of *LDB3* were taken from GenBank (https://www.ncbi.nlm.nih.gov/genbank/) and Protein database (https://www.ncbi.nlm.nih.gov/protein/) for all the *in-silico* analyses. Multiple sequence alignment of LDB3 protein (NP_001073583.1) homologs across different species was performed using the ‘HomoloGene’ feature of NCBI to evaluate the conservation of substituted amino acids in case of non-synonymous variants.

#### 2.3.2 Prediction of pathogenic potential of variants identified in LDB3

The disease-causing potential of non-synonymous variants identified in *LDB3* was predicted using VarCards (http://159.226.67.237/sun/varcards/), which incorporates more than 20 *in silico* predictive algorithms e.g., PolyPhen-2 (http://genetics.bwh.harvard.edu/pph2/), SIFT (http://sift.jcvi. org/www/SIFT_enst_submit.html), Mutation Taster (http://www.mutationtaster.org), I-Mutant (http://folding.biofold.org/i-mutant/i-mutant2.0. html), FATHMM (http://fathmm.biocompute.org.uk/fathmmxf/index.html), VEST3 (https://karchinlab.org/apps/appVest.html), M-CAP (http://bejerano.stanford.edu/mcap/), PROVEAN (Protein Variant Effect Analyzer, http://provean.jcvi.org/index.php) and CADD (https://cadd.gs.washington.edu/).

#### 2.3.3 Effect of non-synonymous variants on secondary-, tertiary structure and physicochemical properties of protein

The secondary structure of the wild-type and mutant LDB3 protein was predicted using the PDBsum (https://www.ebi.ac.uk/thornton-srv/databases/pdbsum/Generate.html) prediction tool. To further, elucidate the effect of non-synonymous variants on the overall protein structure, three-dimensional modelling of LDB3 protein (both wild type and mutants) was performed using Alphafold2 googlecolab (https://colab.research.google.com/github/ <u>sokrypton/ColabFold/blob/main/AlphaFold2.ipynb</u>). For visualizing the differences in the modeled structures across wild-type and mutant LDB3 protein, the two structures were aligned through UCSF Chimera. The Root Mean Square Deviation (RMSD) values were calculated for each of the aligned structures.

The changes in the physico-chemical properties (hydrophobicity, % buried residues, and total β-strand) of LDB3 protein due to non-synonymous variants were computed using the ProtScale (https://web.expasy.org/protscale/) server available at ExPasy portal. The profile for each of the physico-chemical property was obtained and compared across wild-type and mutant LDB3 protein to predict the effect of identified variants.

### 2.4 *In vitro* characterization of the non-synonymous variants

#### 2.4.1 Cloning of wild type *LDB3* and Site Directed Mutagenesis

The wild-type (WT) *LDB3* clone in pEGFP-C1 vector was a kind gift from Prof. Ju Chein’s Lab, UCSD, San Diego. WT-*LDB3* was used as template for preparing mutant constructs of ZASP p.S184I, p.K204R, p.R229C, p.P295T, p.Q402P, p.F465I, p.F496L, p.P606S by site-directed mutagenesis using Quick change II XL site-directed mutagenesis kit (Agilent Technologies Inc., Santa Clara, CA, USA) through mega-primer method as per the manufacturer’s instructions.

#### 2.4.2 Expression of *LDB3* in Cultured Mouse Myoblasts (C2C12 cell line)

Immunostaining was performed to visualize cellular localization and cytoskeletal assembly of WT and mutant *LDB3*. C2C12 cells were transfected with WT and mutant *LDB3* (MUT) for transient expression. The cells were seeded in each well of a six-well culture plate on glass coverslips and grown at 37°C in 5% CO_2_, overnight, in Dulbecco’s modified Eagle’s medium (DMEM, Gibco, Life Technologies Corp., NY, USA), supplemented with 10% Fetal Bovine Serum (FBS, Gibco, Life Technologies Corp., NY, USA). Next day, cells (at 40-45% confluency) were transfected with 500 ng of EGFP-C1-tagged, either WT or MUT-*LDB3* plasmid DNA, using FuGENE 6 transfection reagent (Promega Corp., IN, USA). After 48 hours of transfection, C2C12 cells were washed with 1X PBS twice and immediately fixed with 4% paraformaldehyde (PFA) for 15 minutes, followed by permeabilization with 0.5% Triton X (diluted in 1X PBS). Further, these permeabilized and fixed cells were stained with Phalloidin (10ug/ml; Sigma-Aldrich, MO, USA) and nuclei were stained with DAPI (1 µg/ml), mounted with mounting media and imaging was carried out using Zeiss LSM 510 Meta, Laser Scanning Confocal microscope, further analysed with ZEN 12 and LSM software and assembly of the images were performed using Adobe Photoshop software.

#### 2.4.3 Estimation of protein expression through Western Blotting

The expression level of LDB3 protein in WT versus MUTs was estimated in C2C12 cells, seeded in six-well plate and then transfected with 500 ng each of wild-type and mutant LDB*3* constructs (in EGFP-C1 vector) at 30-40% confluency of cells. Forty-eight hours post-transfection, cell lysates were prepared in RIPA buffer (50 mM Tris-HCl, pH 8.0, 150 mM NaCl, 0.1% Triton X, 0.5% sodium deoxycholate, 0.1% SDS) with 1mM sodium orthovanadate, 1mM NaF, and protease inhibitor tablets (Roche, Basel, Switzerland). After dilution with Laemmli’s sample buffer, the lysates were separated on 10% polyacrylamide SDS gel, and proteins were transferred to PVDF membrane (Bio-Rad Laboratories Inc, CA, US). The PVDF membrane was blocked for 2 hours in 5% dry skimmed milk at room temperature, followed by overnight incubation with anti-GFP antibody (Invitrogen, USA) at 4°C. The blot was washed with PBST (0.1 % Tween-20 diluted in 1X PBS) six times for 5 minutes each. The blot was then incubated with HRP-conjugated goat anti-mouse IgG antibody (Genei, Merck, Germany) followed by washing with PBST (6X, 5 min each) and detected with ECL detection kit (GE Healthcare, IL, USA).

## 3.0 Results

### 3.1 Identification of Genetic Variants in LDB3

Direct DNA sequencing of coding region and exon-intron boundaries of *LDB3/ZASP* in 100 probands by Sanger’s method has revealed a total of 9 non-synonymous, 7 synonymous, and 7 intronic variants. The nine non-synonymous variants are, c.551G>T & c.552C>A (p.S184I), c.577G>A (p.D193N), c.611A>G (p.K204R), c.685C>T (p.R229C), c.883C>A (p.P295T), c.1205A>C (p.Q402P), c.1393T>A (p.F465I), c.1488T>A (p.F496L) and c.1816C>T (p.P606S) (Figure 1). None of these variants is present in 100 ethnic-matched control individuals (200 chromosomes) from the same geographical locality. All these non-synonymous variants have been submitted in ClinVar database, the RCV and SCV numbers have been detailed in Table 1.

**Figure 1.**
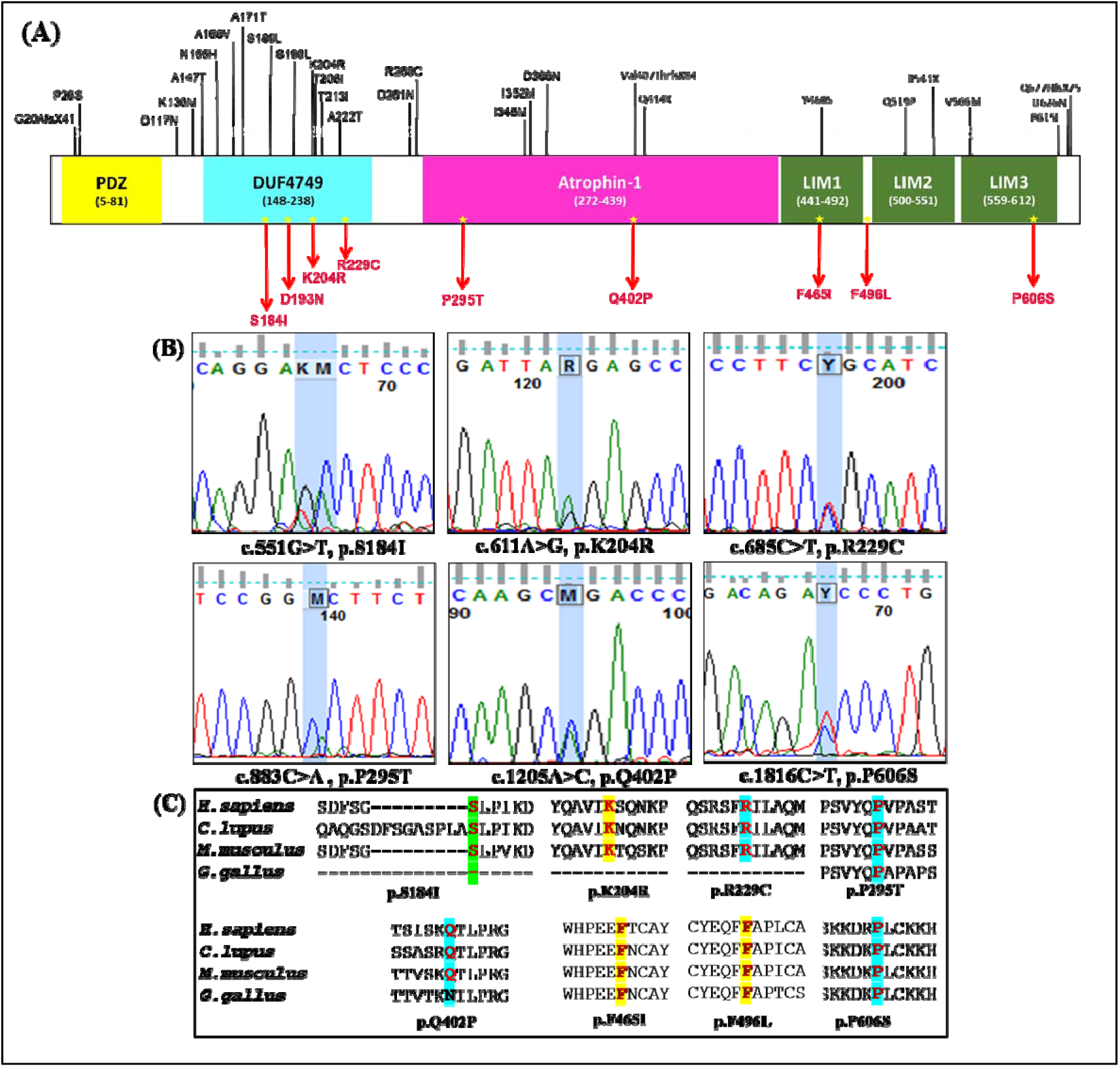
Schematic representation of localization, sequence chromatograms and phylogenetic conservation of LDB3 non-synonymous variants. **(A)** LDB3 protein -showing a PDZ domain, domain of unknown function 4749 (DUF4749), atrophin-1 domain and three LIM domains viz., LIM1, LIM2, LIM3. Position of identified non-synonymous variants (present study) are marked in red color while previously published LDB3 variants in DCM and other forms of cardiomyopathies are marked in black color. **(B)** Sequence chromatogram of mutant genetic variants. **(C**)Phylogenetic conservation of LDB3 protein across different species showing substituted amino acid residues highlighted in different colors.

**Table 1.** List of non-synonymous variants identified in *LDB3* in DCM cases showing location of the identified variants, minor allele frequency, and clinical characteristics.

| Nucleotide Change | Domain | No. of Case (s) (MAF) | No. of Control (s) | rsID/RCV ID (MAF) | INDEX-db | IndiGenomes | LVEF (%) | LVIDd (mm) |
| --- | --- | --- | --- | --- | --- | --- | --- | --- |
| **c.551G>T & c.552C>A (p.S184I)** | DUF4749 | 1/100 (0.005) | 0/100 | **Novel**<br>(RCV001293335.1 & RCV001293336.1) | NR | NR | 35 | 65 |
| **c.577G>A (p.D193N)** | DUF4749 | 1/100 (0.005) | 0/100 | **Novel**<br>(RCV001293337.1) | NR | NR | 56 | 44 |
| **c.611A>G (p.K204R)** | DUF4749 | 1/100 (0.005) | 0/100 | rs34423165 (0.0156) | 0.02 | 0.0064 | 62 | 77 |
| **c.685C>T (p.R229C)** | DUF4749 | 1/100 (0.005) | 0/100 | rs397517226 (NR) | NR | 0.0005 | 70 | 65 |
| **c.883C>A (p.P295T)** | Atrophin-1 | 1/100 (0.005) | 0/100 | **Novel**<br>(RCV001293340.1) | NR | NR | 62 | 77 |
| **c.1205A>C (p.Q402P)** | Atrophin-1 | 1/100 (0.005) | 0/100 | rs138951890 (0.0007) | NR | NR | 35 | 65 |
| **c.1393T>A (p.F465I)** | LIM1 | 1/100 (0.005) | 0/100 | **Novel**<br>(RCV001293342.1) | NR | NR | 35 | 65 |
| **c.1488T>A (p.F496L)** | - | 1/100 (0.005) | 0/100 | **Novel**<br>(RCV001293343.1) | NR | NR | 20 | 78 |
| **c.1816C>T (p.P606S)** | LIM3 | 1/100 (0.005) | 0/100 | **Novel**<br>(RCV001293344.1) | NR | NR | 33 | 57.6 |

Footnotes: Nomenclature refers to *LDB3* reference sequence (NM\_001080114.2). For cDNA positions, +1 refers to the base A of first codon 'ATG'; for protein (NP\_001073583.1) positions +1 refers to amino acid coded by first codon 'ATG'. DUF4749: domain of unknown function, MAF: Minor Allele Frequency, NR: not reported, rsID: reference sequence ID, LVEF=Left ventricular ejection fraction, LVIDd= Left ventricular internal end diastolic diameter, RCV: clinical variation database Reference number of submitted novel variants

Out of these nine non-synonymous variants, six are novel c.551G>T & c.552C>A (p.S184I), c.577G>A (D193N), c.883C>A (p.P295T), c.1393T>A (p.F465I), c.1488T>A (p.F496L) and c.1816C>T (p.P606S). Interestingly, one of the variant p.S184I possess two consecutive base substitutions (c.551G>T & c.552C>A) in the same codon. Here the two successive base changes resulted in a single amino acid substitution at 184^th^ position, where the amino acid serine is replaced by isoleucine. Hence the change referred as p.S184I.

In addition to these, three other variants viz., c.611A>G (p.K204R), c.685C>T (p.R229C) and c.1205A>C (p.Q402P), are already reported in the gnomAD database (Table 1). The minor allele frequencies (MAF) of all the novel non-synonymous variants identified in our study are found to be <0.005. MAF of two of the already reported variants, viz. p.K204R and p.Q402P, was 0.0156 and 0.0007, respectively, in gnomAD, whereas the MAF of another known variant, p.R229C, was not reported in the gnomAD database. The mining of INDEX-db and IndiGen*omes* databases, which encompasses variants identified in healthy individuals (controls) from the Indian population, indicated that p.K204R has been reported in both INDEX-db (MAF=0.02) and IndiGen*omes* (MAF=0.0064) databases. The variant p.R229C has been reported in IndiGen*omes* (MAF=0.0005) but not in INDEX-db. No other variants are found in both these databases. MAF of nine non-synonymous variants and the clinical phenotype (left ventricular) of patients are shown in Table 1.

The clinical characteristics of patients harboring these variants are severely impaired. The variants p.S184I, p.Q402P, and p.F465I are identified in a single patient who is a 3.5-year-old female with severely reduced ejection fraction (LVEF) of 35% and enlarged end-diastolic diameter (LVIDd) of 65mm. Similarly, two other variants, p.K204R and p.P295T, are also identified in a 4-month-old male patient with a normal LVEF of 62% however, severely enlarged LVIDd of 77 mm. The ejection fraction (LVEF) and left ventricular end-diastolic diameter (LVIDd) are listed in Table 1. The variant, c.577G>A (p.D193N), was a novel variation, not reported in any of the control databases, including the Indian control-database (IndiGen*omes and* INDEX-db). Three variants, p.S184I, p.K204R, and p.R229C are present in the DUF4749 domain, the other two, p.P295T and p.Q402P, in the Atrophin-1 domain, p.F465I in the LIM1 domain, and p.P606S in the LIM3 domain, while the variant, p.F496L, is positioned between 1^st^ and 2^nd^ LIM domains. All 3 variants present in the LIM domain have severe phenotype with reduced ejection fraction and enlarged LVIDd. Among these, p.F496L exhibits most severe phenotype with a very low LVEF of 20% and LVIDd of 78 mm, followed by LIM3 domain variant p.P606S (LVEF= 33% and LVIDd=57.6mm).

Apart from the nine non-synonymous variants, 7 synonymous (Table 3) and 7 intronic variants (Supplementary Table 1) have also been identified in *LDB3.* The synonymous variants are c.273G>A (p.T91=), c.456G>A (p.A152=), c.504T>C (p.D168=), c.546T>C (p.S182=), c.837GCC>GCT (p.A279=), c.1041C>A (p.S347=) and c.1074C>T (p.A358=) (Figure 2A). None of these synonymous variants could be detected in our control cohort from the same geographic location. However, all of these are reported in dbSNP, albeit the MAF of most of these are <0.01 except for two, i.e., p.D168= (MAF=0.039) and p.A358= (MAF=0.032) (see Table 3). The seven intronic variants were c.322-50G>C, c.548+53A>C, c.718+47G>C, c.718+81G>A, c.755+11G>A, c.755+86G>A, and c.*30C>G (Figure 2B).

**Figure 2:**
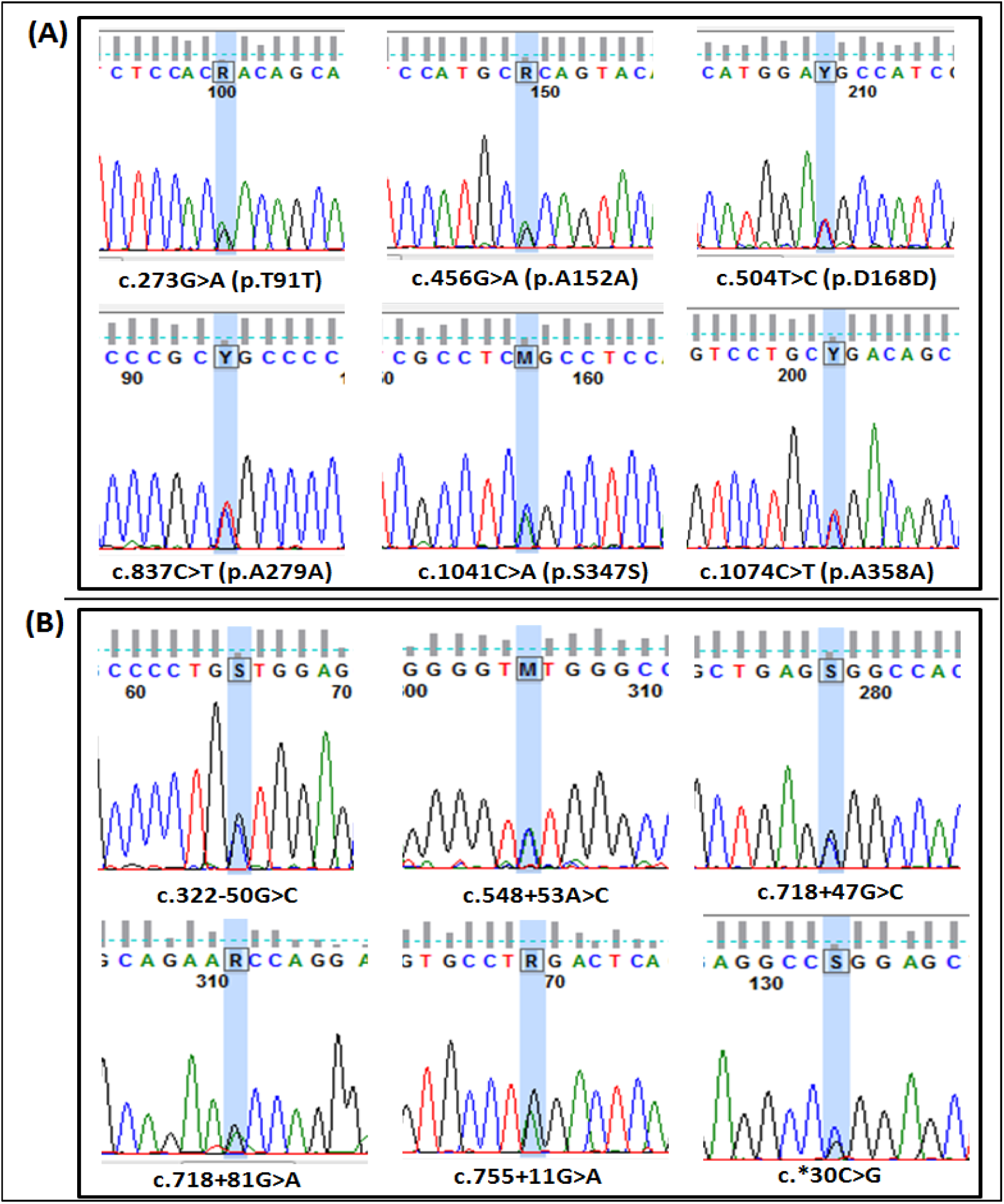
Sequence chromatogram of (A) synonymous variants and (B)intronic variants identified in LDB3.

**Table 2:** Pathogenic potential of the non-synonymous variants, predicted using VarCards.

| VARIANT | PATHOGENICITY (*in silico*) |  |  |  |  |  |  |  |  |
| --- | --- | --- | --- | --- | --- | --- | --- | --- | --- |
|  | Polyphen2H<br>DIV<br>(Cut off=0.5) | Mutation<br>Taster | SIFT<br>(Cut<br>off=0.05) | VEST3<br>(Cut off=0.5) | FATHMM<br>(Cut<br>off=0.05) | M_CAP | PROVEAN | CADD | I-Mutant<br>(DDG in<br>Kcal/mol) |
| **c.551G>T & c.552C>A**<br>**(P.S184I)** | Probably<br>Damaging<br>(0.958) | Disease<br>Causing (1) | Damaging<br>(0.021) | Tolerable<br>(0.435) | Tolerable<br>(0.55) | Damaging<br>(0.145) | Damaging<br>(-2.55) | Tolerable<br>(10.07) | Increase<br>Stability |
| **c.611A>G (P.K204R)** | Probably<br>Damaging<br>(0.998) | Disease<br>Causing (1) | Tolerable<br>(0.501) | Damaging<br>(0.712) | Tolerable<br>(0.69) | - | Tolerable<br>(-0.97) | Damaging<br>(23.2) | Decrease<br>Stability |
| **c.685C>T (P.R229C)** | Probably<br>Damaging<br>(1) | Disease<br>Causing (1) | Damaging<br>(0.001) | Damaging<br>(0.915) | Tolerable<br>(0.18) | Damaging<br>(0.034) | Damaging<br>(-5.48) | Damaging<br>(35) | Increase<br>Stability |
| **c.883C>A (P.P295T)** | Probably<br>Damaging<br>(1) | Disease<br>Causing (1) | Tolerable<br>(0.285) | Damaging<br>(0.808) | Damaging (-<br>1.7) | Damaging<br>(0.038) | Tolerable<br>(-2.14) | Damaging<br>(23) | Decrease<br>Stability |
| **c.1205A>C (P.Q402P)** | Probably<br>Damaging<br>(0.986) | Disease<br>Causing (1) | Tolerable<br>(0.297) | Damaging<br>(0.542) | Tolerable<br>(0.78) | Damaging<br>(0.077) | Tolerable<br>(-1.77) | Tolerable<br>(15.53) | Decrease<br>Stability |
| **c.1393T>A (P.F465I)** | Probably<br>Damaging<br>(1) | Disease<br>Causing (1) | Damaging<br>(0.001) | Damaging<br>(0.933) | Damaging (-<br>3.27) | Damaging<br>(0.221) | Damaging<br>(-4.3) | Damaging<br>(31) | Decrease<br>Stability |
| **c.1488T>A (P.F496L)** | Probably<br>Damaging<br>(0.999) | Disease<br>Causing (1) | Damaging<br>(0.019) | Damaging (0.84) | Damaging (-<br>2.42) | Damaging<br>(0.122) | Damaging<br>(-3.95) | Damaging<br>(25.7) | Decrease<br>Stability |
| **c.1816C>T (P.P606S)** | Probably<br>Damaging<br>(1) | Disease<br>Causing (1) | Damaging<br>(0.002) | Damaging<br>(0.898) | Damaging (-<br>2.26) | Damaging<br>(0.196) | Damaging<br>(-5.27) | Damaging<br>(33) | Decrease<br>Stability |

**Table 3:** List of synonymous variants identified in *LDB3* and their impact on RNA and protein expression.

| Genetic variant | No. of Case (s) (MAF) | No. of Control (s) | INDEX-db | IndiGenomes | rsID/MAF gnomAD | MFold (mRNA Folding and stability) | δRSCU (Codon preference during translation with respect to *LDB3* CDS) | CUF (Codon preference during translation with respect to *LDB3* CDS) |
| --- | --- | --- | --- | --- | --- | --- | --- | --- |
| c.273G>A<br>p.T91= | 6/100<br>(0.03) | 0/100 | 0.0285 | 0.0249 | rs45613039<br>(0.002) | Altered, Decreased | Mutant Codon | Mutant Codon |
| c.456G>A<br>p.A152= | 1/100<br>(0.005) | 0/100 | 0.0028 | 0.0029 | rs371708921<br>(0.0004) | Not altered, Increased | Mutant Codon | Mutant Codon |
| c.504T>C<br>p.D168= | 2/100<br>(0.01) | 0/100 | 0.0273 | NR | rs76615432<br>(0.039) | Altered, Decreased | Mutant Codon | Mutant Codon |
| c.546T>C<br>p.S182= | 1/100<br>(0.005) | 0/100 | NR | 0.0088 | rs71473272<br>(0.005) | Altered, Decreased | Mutant Codon | Mutant Codon |
| c.837C>T<br>p.A279= | 1/100<br>(0.005) | 0/100 | NR | NR | rs768844187<br>(NR) | Not altered, Decreased | Wild-type Codon | Wild-type Codon |
| c.1041C>A<br>p.S347= | 3/100<br>(0.015) | 0/100 | NR | 0.0523 | rs45555240<br>(NR) | Altered, Increased | Wild-type Codon | Wild-type Codon |
| c.1074C>T<br>p.A358= | 1/100<br>(0.005) | 0/100 | NR | 0.0020 | rs45459491<br>(0.032) | Altered, Increased | Wild-type Codon | Wild-type Codon |

Three of the intronic variants are already reported while four are novel. However, these are far off the splice sites; therefore, *in-vitro* functional characterization for none of the intronic or synonymous variants *Homo sapiens* (NP_001073583.1), *Canis lupus* (XP_005619166.1), *Mus musculus* (NP_001034161.1), and *Gallus gallus* (XP_004942002.1) has revealed that the majority of the substituted amino acids are conserved across mammalian species.

The N-terminal domain of LDB3 (1-68 amino acids; 82-92 amino acids) is unknown in case of Gallus. However, most of the C-terminal variants, namely p.P295T, p.F465I, p.F496L, p.P606S, show conservation with *G.gallus* except p.Q402P. The conserved region showing the substituted amino acids, along with flanking amino acid sequences, is shown in Figure 1C.

### 3.3 Pathogenic potential of LDB3 nonsynonymous variants

We have used *VarCards* (http://159.226.67.237/sun/varcards/) for prediction of pathogenic potential of the non-synonymous variants. Out of 23 tools, that *VarCards* encompasses, the variant p.Q402P is predicted to be deleterious by 10 tools, while p.S184I by 11, p.K204R by 13, p.F496L by 17, p.P295T by 18, p.R229C by 19, p.F465I and p.P606S by 21 tools. The predictions of 9 important tools have been shown in Table 2.

### 3.4 Prediction of structural changes (secondary and tertiary) in LDB3 protein

Prediction of secondary structure using PDBsum has demonstrated remarkable changes in the secondary structure of mutant LDB3 protein due to all the non-synonymous variants including insertion of alpha-helix, beta-sheets and hairpins, removal of helix etc. The detailed changes in the secondary structure of the protein have been listed in Supplementary figure 1 and supplementary table 2) which clearly indicates that highest level of secondary structure changes were observed in the variants p.P295T and p.S184I (Supplementary Figure 1), followed by p.F496L, p.F465I, p.Q402P, p.P606S, p.R229C,p.D193N and p.P295T.

Three-dimensional protein structure of wild-type and mutant LDB3 protein has been modelled and aligned to examine the structural differences. We have observed that due to all the variants there are significant changes in the structure of the mutant protein as evident by the root mean square deviation (RMSD) values. The RMSD values (in Å) of variants p.S184I, p.D193N, p.K204R, p.R229C, p.P295T, p.Q402P, p.F465I, p.F496L and p.P606S are estimated to be 5.491, 1.168, 4.44, 6.75, 1.021, 0.941, 3.328, 0.766, and 0.916 respectively (Figure 3). Furthermore, the changes in physico-chemical properties of LDB3 protein due to non-synonymous variants revealed the total beta strand also shown significant changes due to all the variants. The profile for hydrophobicity and total beta strand of variant, p.S184I has been shown in Supplementary Figure 2.

**Figure 3:**
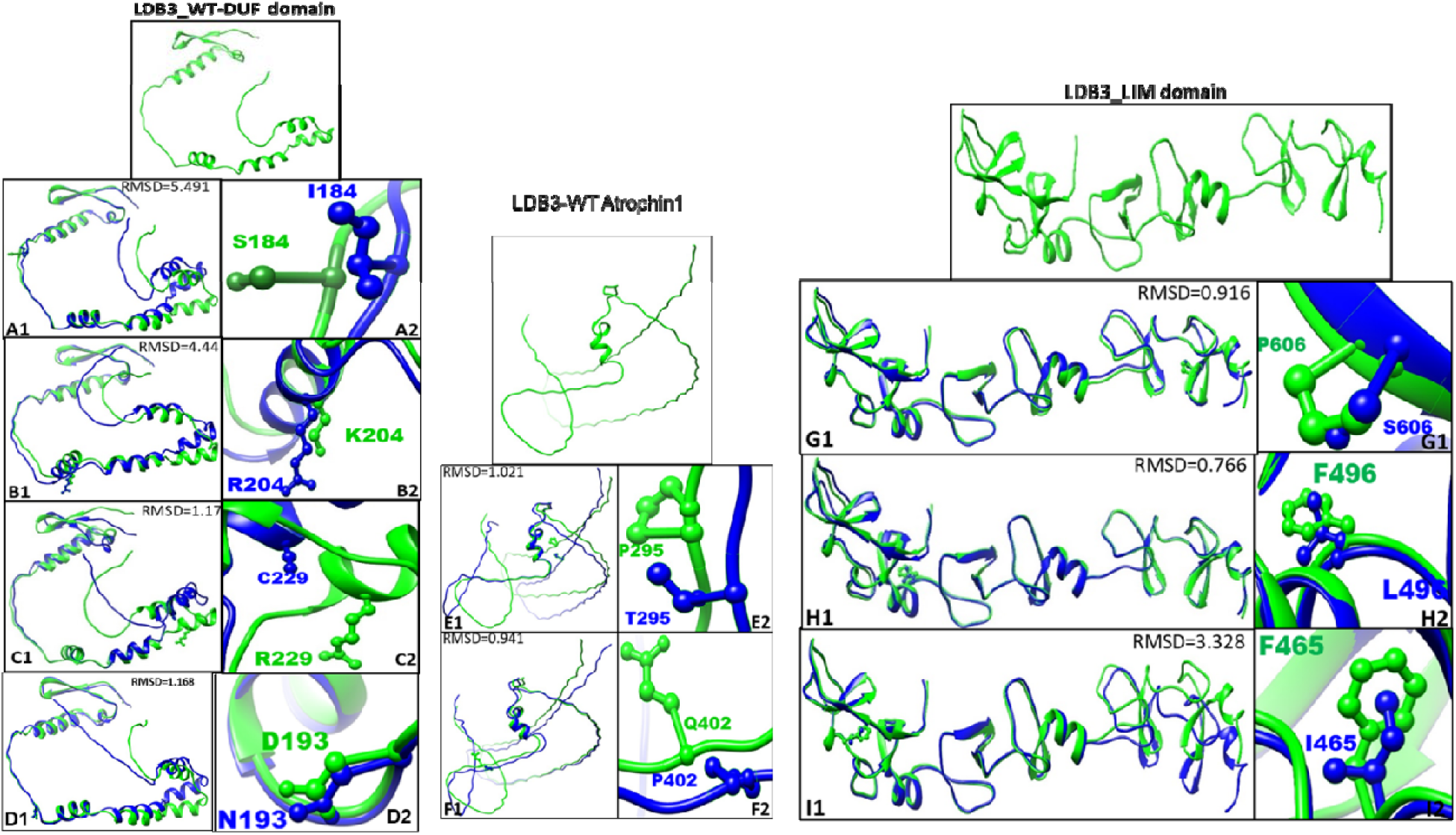
Structural models of the human LDB3 protein were generated in a domain-wise manner, with the DUF, Atrophin1 and LIM domains predicted using AlphaFold2, which provides high-confidence predictions based on deep-learning-assisted structure inference. Separate models were constructed for both the wild-type and mutant forms of each domain to investigate the potential structural impact of the identified amino acid substitution(s). The resulting models were subsequently superimposed to visualize local and global conformational variations between the wild-type and mutant structures (A1-I2). The root mean square deviation (RMSD) values, representing the extent of atomic displacement and structural divergence, are displayed within each figure panel, highlighting regions most affected by mutation-induced three-dimensional structural alterations.

### 3.5 Impact of non-synonymous variants on the expression and localization of LDB3 protein

The expression of wild-type and mutant LDB3 protein in C2C12 cells has been evaluated by Western blotting. The results have shown significant differences in the level of protein expression. The expression of LDB3 protein is down-regulated in variants p.S184I (1.58 fold, p<0.04), p.F465I (1.35 fold, p<0.02), p.F496L (1.61 fold, p<0.04) and p.P606S (1.25 fold, p<0.04) as compared to wild type (Figure 4) whereas it was upregulated in variant p.K204R (1.25 fold, p<0.02). In other variants, there was no significant difference in the expression.

**Figure 4:**
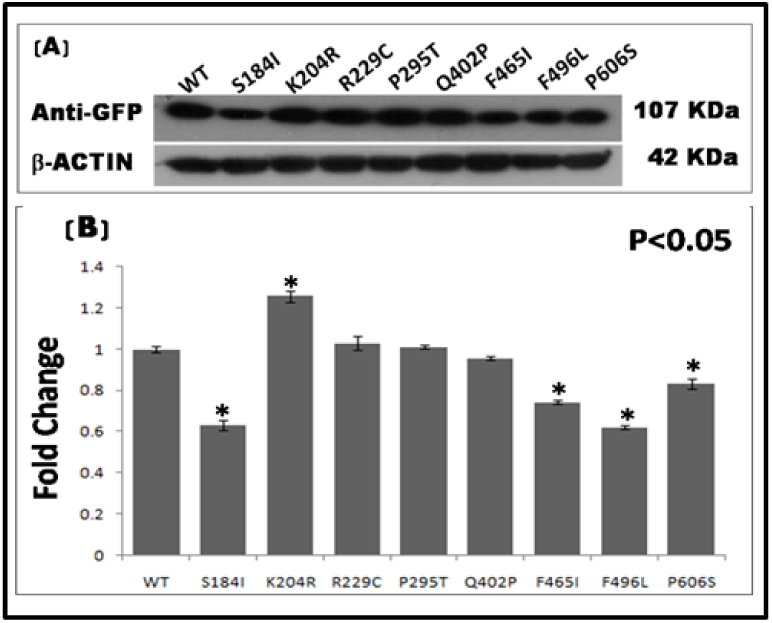
Expression of wild-type and mutant LDB3 proteins. (A) Western blotting (B) The bar chart representing mean fold change in the expression of wild-type and mutant proteins, calculated from three independent experiments. P value <0.05 is considered statistically significant.

The cellular localization of wild-type and mutant LDB3 has also been studied, by immunofluorescence, which demonstrate its localization along the Z-disc in both wild-type and mutants. The variant p.K204R has shown over-expression of LBD3. Interestingly, blebs of cytoplasmic aggregation of the protein have been recorded. The variant p.R229C also over-expressed the mutant protein. In case of variant, p.P295T, Z-disc was severely disrupted (Figure 5). We also noted prominent disruption of actin-based cytoskeleton in variants p.K204R, p.R229C, p.P295T, p.Q402P, p.F465I, p.F496L.

**Figure 5:**
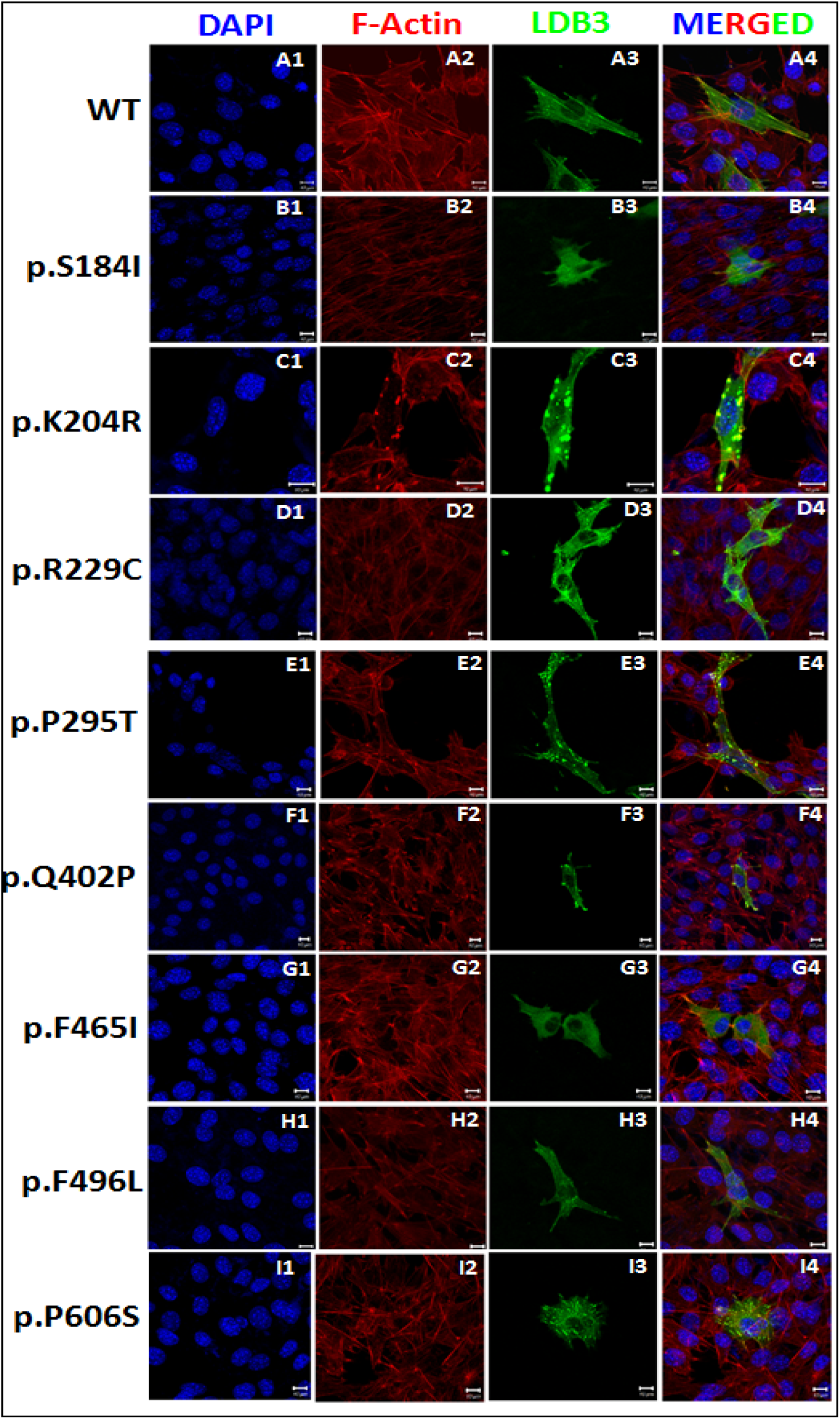
Expression of LDB3 in C2C12 cells. (A1-I4) Immunofluorescence of C2C12 cells showing expression of LDB3 in wild-type, p.S184I, p.K204R, p.R229C, p.P295T, p.Q402P, p.F465I, p.F496L, p.P606S

### 3.6 Effect of synonymous variants on mRNA folding

A comparative analysis of wild-type and mutant mRNA has revealed striking changes in the conformation and rearrangement of nucleotides in case of variants c.273G>A (p.T91=), c.504T>C (p.D168=), c.546T>C (p.S182=), c.1041C>A (p.S347=) and c.1074C>T (p.A358=) (Figure 4.9). However, in case of variants c.456G>A (p.A152=) and c.837GCC>GCT (p.A279=) we observe no differences in the conformation of mRNA.

In case c.273G>A (p.T91=), the wild-type allele "G" (red circled) is located in a multi-branch loop at the base of short hairpin while the mutated allele "A" is located at a different place in the loop (Figure 4.9A). First bulge loop of the wild-type RNA is lost in the mutant mRNA. In c.504T>C (p.D168=) mRNA folding showed noticeable differences between wild-type and mutant mRNA (Figure 4.9). Wild-type mRNA comprises of two internal loops, two small buldge loops and a terminal loop while the mutant mRNA possess three internal loops, a buldge loop and a terminal loop. Besides, one more small helix is also present in the mutant mRNA. Wild type allele is located in the helix while mutated allele is located in an internal loop. In case of c.546T>C (p.S182=), the wild-type mRNA has only single internal loop while the mutated mRNA has two internal loops (Figure 4.9C). In case of c.1041C>A (p.S347=), the multi-branch loop contains 4 branches (helices or hairpin) while mutant mRNA has only three helices (Figure 4.9D). The wild-type allele is located in a hairpin loop in the wild-type mRNA folding while mutated allele is a part of a hairpin. In wild-type mRNA, one of the helix has an internal buldge-loop which has been replaced by an internal loop in the secondary structure of mutated RNA. The multi-branch loop of wild type mRNA contain two hairpins, one of which possesse an internal loop while in mutant (c.1074C>T, p.A358=), the multi-branch loop possesse three hairpins. Here, wild-type allele "C" is part of a short hairpin while mutant allele "T" is the part of a hairpin loop (Figure 6).

**Figure 6:**
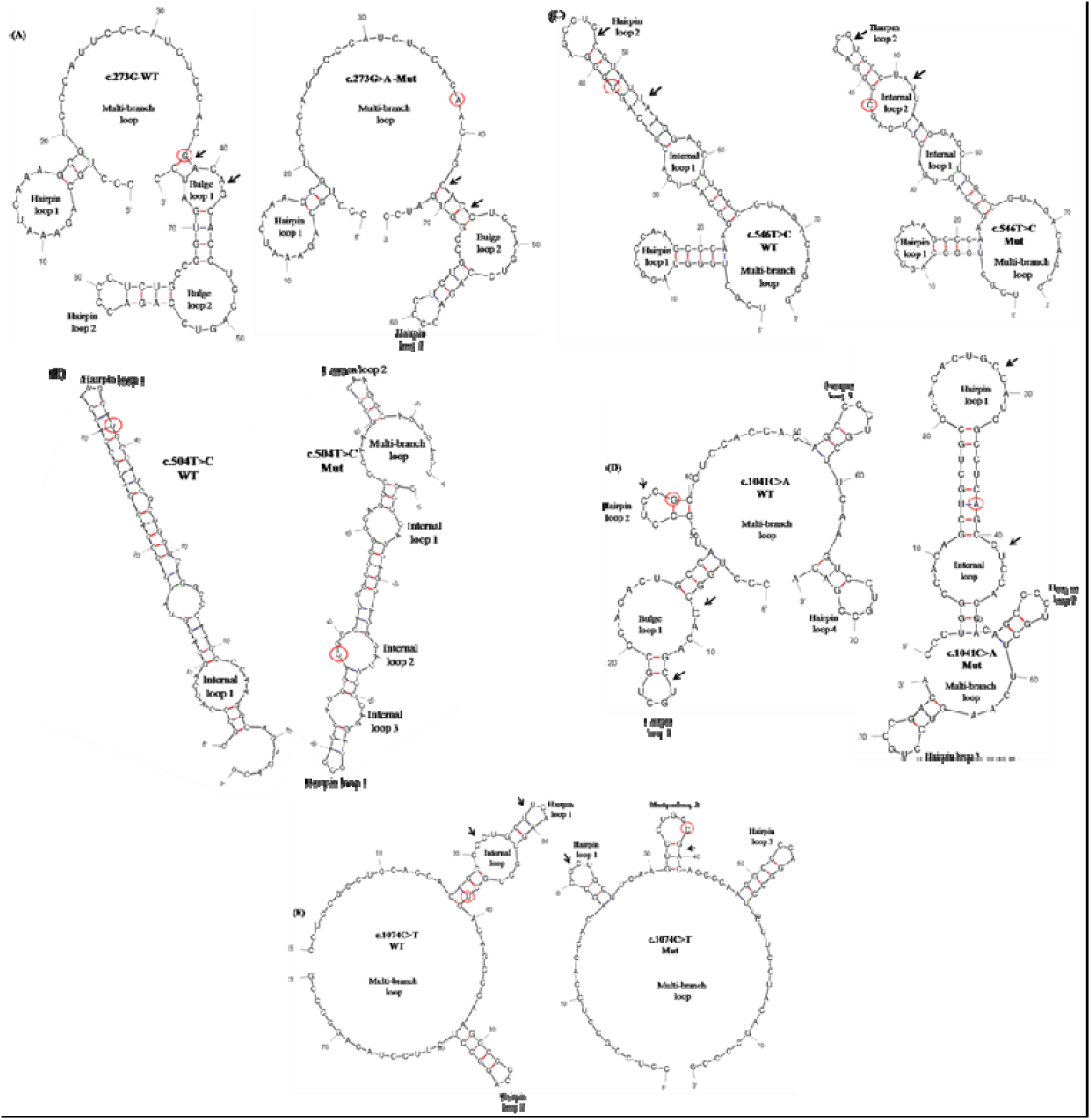
Schematic representation of mRNA structure, and stability predicted using Mfold. Picture showing mRNA structures of 75 nucleotide long fragment for (A) c.273G>A, (B) c.504T>C, (C) c.546T>C, (D) c.1041C>A and (E) c.1074C>T. The wild-type and mutant structures are placed side by side for comparison. The WT and the mutated bases are indicated with red circles and the regions harboring structural changes in the mutant mRNA due to variant of interest are indicated with arrows.

We have also analyzed the stability of mutant mRNA structures compared to wild-type from the calculated differences in the minimum free energy values. The differences in the minimum free energy values (δδG) of variants c.273G>A (p.T91=), c.456G>A (p.A152=), c.504T>C (p.D168=), c.546T>C (p.S182=), c.837GCC>GCT (p.A279=), c.1041C>A (p.S347=) and c.1074C>T (p.A358=) for 75 long nucleotide structure were 0.3, 0, 1.5, 1.5, 0.3, -0.7 and -0.1 respectively. Likewise, these values for all the aforementioned variants in case of 151 nucleotide structure are 1.9, 0.7, 2.2, 1.5, 0.3, -0.7 and 0 respectively. The more positive is the δδG value the less stable is the mutant mRNA. As shown in Figure 7, it is evident that the stability of predicted mutant mRNA structure decreased in case of variants c.273G>A (p.T91=), c.504T>C (p.D168=), c.546T>C (p.S182=), c.837C>T (p.A279=) irrespective of nucleotide length, while in case of variant c.456G>A (p.A152=) there is no difference in the stability compared to wild-type for 75 nucleotide structure but in this variant also stability of predicted mRNA structure decreased for 151 nucleotide structure. However, in case of variant c.1041C>A (p.S347=) the stability of predicted mRNA increased irrespective of nucleotide length. Similar analysis has been performed for variant c.1074C>T (p.A358=) which has revealed that there is no difference in the stability for 75 nucleotide long predicted structure but for 151 nucleotide structure the stability has increased.

**Figure 7:**
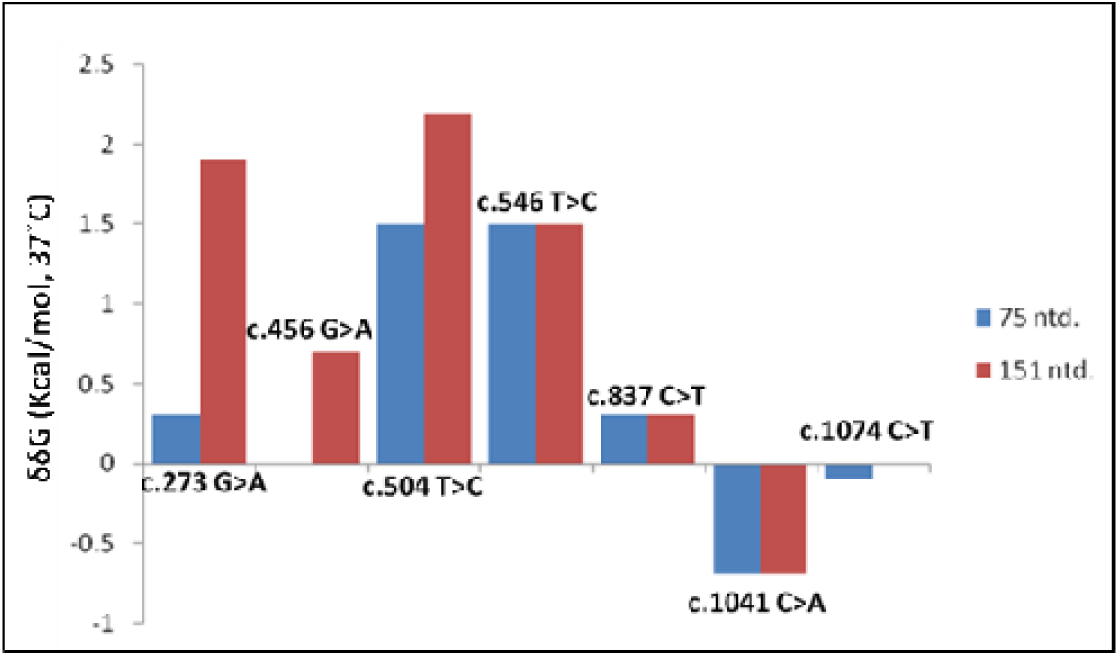
Representation of Mfold δδG values of LDB3 synonymous variants. RNA fragments of 75 and 151 nucleotides in length were used as input sequence in the Mfold server, with default server settings. The most stable structures (lowest δG) for variants and wild-type LDB3 were selected. The δδG (δGMutant – δGWild-type) was calculated and plotted for each of the variants. A more positive δδG value is indicative of less stable mutated mRNA compared to wild-type mRNA

### 3.7 Computational analysis of Relative Synonymous Codon Usage (RSCU) and Codon Usage Frequencies of LDB3 synonymous variants

The RSCU values were calculated for each of the synonymous codon and the corresponding wild-type codon. A comparison was made for variant versus wild-type. The δRSCU values of the synonymous variants, both with respect to *LDB3* gene specific codon usage and Human genome specific codon usage have been listed in Table 4 and also represented in Figure 8.

**Figure 8.**
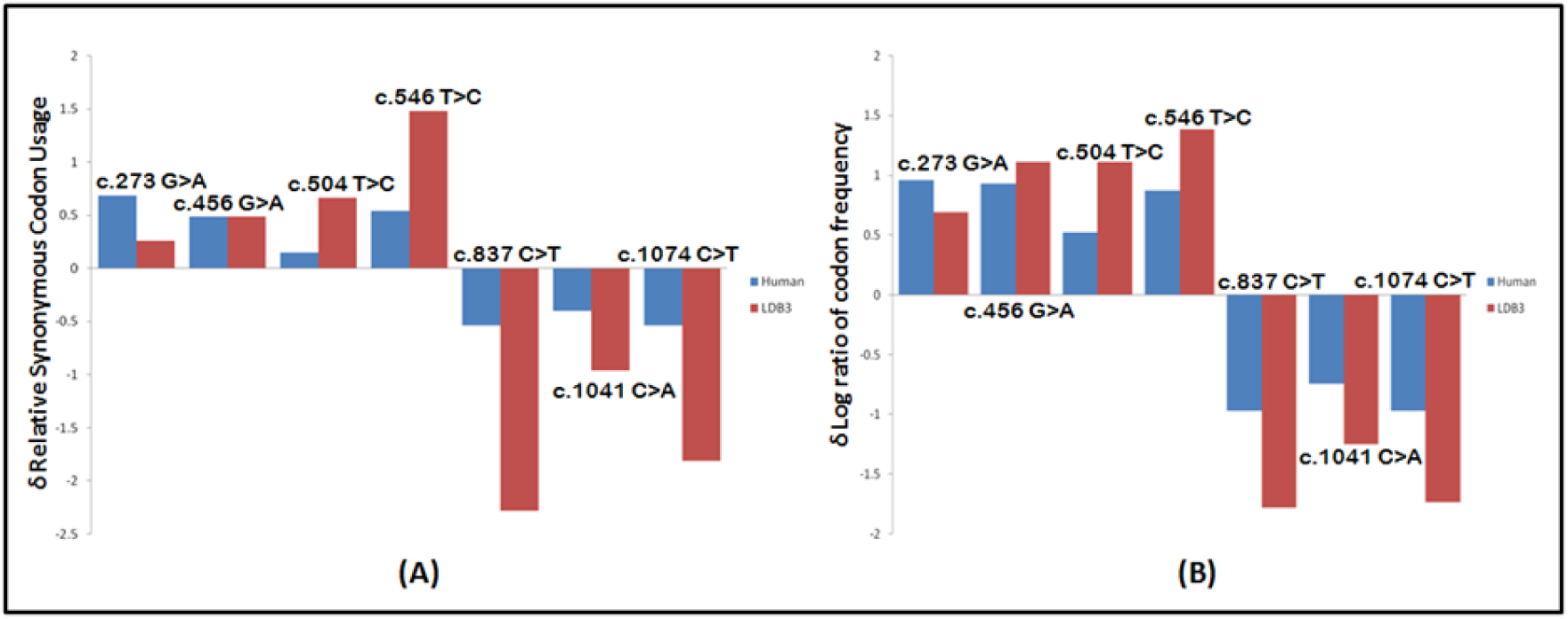
Graphical representation of the codon frequency changes due to synonymous variants in LDB3 (A) The Relative Synonymous Codon Usage (RSCU) method (B) The Log ratio method. RSCU frequency of wild-type and mutant codons were calculated using frequencies derived from the human genome and LDB3. Change in RSCU (δRSCU = RSCU Mutant – RSCU Wild-type) and Log Ratio (δLog ratio = Log ratio Mutant – Log ratio Wild-type) were plotted. A more negative δRSCU or δLog ratio value suggests that the wild-type codon is more common when compared to mutant codon

**Table 4.**
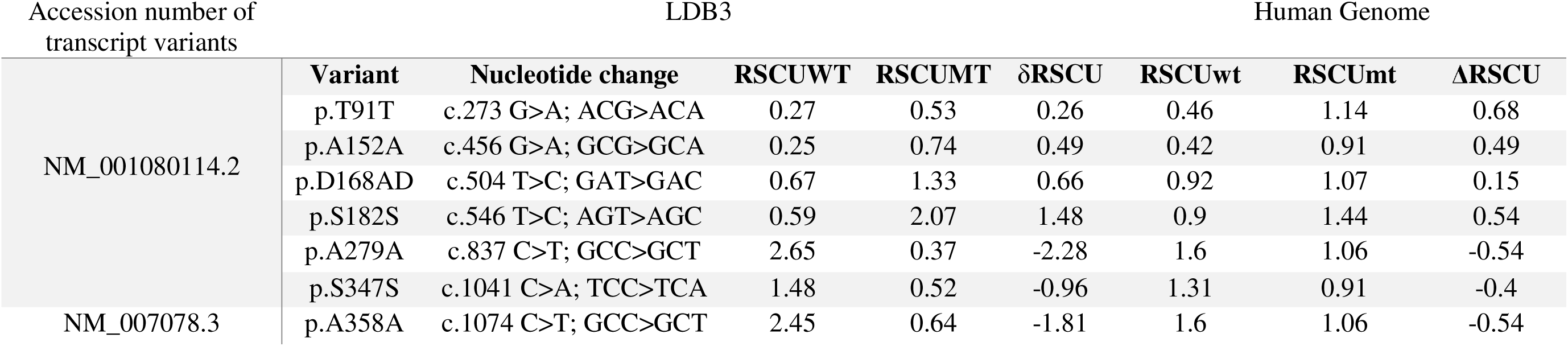
Calculation of relative synonymous codon usage (RSCU) and change in relative synonymous codon usage (δRSCU) with respect to both *LDB3* and Human Genome.

For variants c.273G>A (p.T91=), c.456G>A (p.A152=), c.504T>C (p.D168=), c.546T>C (p.S182=) δRSCU values are 0.26, 0.49, 0.66 and 1.48 respectively whereas for variants c.837GCC>GCT (p.A279=), c.1041C>A (p.S347=) and c.1074C>T (p.A358=), the values are -2.28, -0.96 and -1.81 respectively with respect to *LDB3* gene specific codon usage. These values suggest that in case of variants c.273G>A (p.T91=), c.456G>A (p.A152=), c.504T>C (p.D168=), c.546T>C (p.S182=), mutant codon is more frequently used than wild-type codon whereas in case of variants c.837C>T (p.A279A), c.1041C>A (p.S347S) and c.1074C>T (p.A358A) wild-type codon is more commonly used when compared to mutant codon. The δRSCU values were also calculated for all the seven synonymous variants using the human genome codon usage table and we obtained similar results. The log-ratio for codon usage frequency has been also calculated for all the seven synonymous variants from the *LDB3* codon usage and human genome codon usage database and the results were similar to RSCU.

## 4. Discussion

The LIM domain binding protein 3 (LDB3) is a member of PDZ-LIM domain binding family of protein, that plays a critical role in maintaining the structural integrity of Z-disc in both cardiac and skeletal muscle. It interacts with number of sarcomeric proteins such as ACTN2, Myotilin (MYOT), Calsarsin, Nebulin (NEB), α-Actin (ACTC1), Protein kinase C (PKC) (Zhou *et al*. 1999; von Nandelstadh *et al*. 2009; Arimura *et al*. 2004; Lin *et al*. 2014). It not only helps in maintaining structural architecture of sarcomeric Z-disc but also participate in force generation and transmission during muscle contraction. Thus, LBD3 act as a key mediator in mechanosensing and signaling.

In the present study, we have screened all the sixteen coding exons in LDB3 in a cohort of 100 DCM cases and reported nine non-synonymous (6 novel and 3 previously reported) rare variants in 6 patients. Interestingly, 3 mutations are detected in one patient and 2 mutations in another patient. Rest four mutations are observed each one in four different patients. In addition, we also identified 7 synonymous changes in the protein coding region in 15 sporadic cases, which are not detected in our control cohort, however several of these are reported in Indian control database (IndiGen*omes*) in low frequency (< 0.01), except for two of these (p.T91= in 6 cases and p.S347= in 3 cases) showing higher MAF (0.02 and 0.05 respectively). The overall mutation frequency of *LDB3* in our study (Indian population) appears to be high (21%) of the cohort compared to other published studies, i.e. 6 % in Vatta *et al*. 2003 (American), 2.5% in Theis et al., 2006 (European) and 1.04 % in Arimura et al., 2004 (Japanese).

Here, we report nine non-synonymous variants, of which four (p.S184I, p. D193N, p.K204R and p.R229C) are localized to the DUF4749 domain, 2 (p.P295T and p.Q402P) in Atrophin-1 domain, while one each in LIM1 domain (p.F465I) and LIM3 domain (p.P606S) and one more lies in between LIM1 and LIM2 domain. Extensive survey of literature suggested that a total of 29 mutations have so far been recorded (see supplementary table 3). Out of these 29 mutations, 9 are in DUF4749 domain which contribute 31% of total. We have detected 44.44% (4 out of 9) of mutation in the same domain in our study cohort, which is even higher than other studies. Although function of this domain is not known precisely so far, high frequency of mutations harboured in this domain strongly suggests that this domain has possible functional significance, which is more likely to be associated with disease phenotype. Previously, in another study, Vatta et al., 2003 have also detected two mutations i.e., S196L and T213I, in the same domain. The mutation, S196L has been shown strong disease association in a three-generation family. Further in a transgenic mouse model study, of S196L-TG mice exhibited severe dilation of heart with reduced ejection fraction and fractional shortening at 10 months. Histological study of these 10 months-old mice has also shown LV dilation and altered cyto-architecture. Electron microscopy of myocardium has also clearly demonstrated severe sarcomere disarrangement and altered Z-band structure (Li et al., 2010). Furthermore, isolated cardiomyocytes from 10-month-old TG-S196L mice have shown impaired contractile function as well. This study also claims the interaction of sodium and calcium channels with this domain. Taken together all these evidences, we have reasoned that this domain is likely functionally significant. Although no other mutation in this domain, has so far been thoroughly characterized, the ample evidences from mutant S196L strongly support the functional role of this domain. In the present study, we have identified four non-synonymous variations, namely p.S184I, p.D193N, p.K204R and p.R229C. Although the variant p.D193N is a novel variation and not reported in any of the control databases including Indian population databases, functional characterization could not be performed due to some technical limitations. However, *in vitro* functional characterization of three other variations (p.S184I, p.K204R and p.R229C) has demonstrated altered protein expression and disruption (disarrayed) of actin cytoskeleton. Moreover, *in-silico* secondary and tertiary structure prediction of the mutant proteins have shown remarkable changes in both secondary (Supplementary Table 2, supplementary figure No.1) and tertiary structure (Figure 3). Superimposed confirmation of mutants (p.S184I, p.D193N, p.K204R, p.R229C) over wild-type have clearly shown the distorted mutant protein structure. The root Mean square deviation (RMSD) values (5.491, 1.168, 4.44, and 6.75 respectively) also confirm the deleterious nature of these mutations.

Interestingly, immuno-cytochemistry of the mutant p.K204R, illustrate high level of protein aggregation in the periphery of the cells along with severe destruction of actin cytoskeleton. Similar aggregate formation was evident in Drosophila muscle, where mutant LDB3/ ZASP oligomerise to form pathological aggregates of smaller isoform of LDB3/ ZASP lacking the LIM domain (Gonzalez-Morales; 2019). In human large aggregates of Z-disc proteins are seen in patient suffering from myopathies. Murphy and Young (2015) and Kelly et al., (2016) have described such aggregate formation by any of the Z-disc associated LIM domain proteins, such as α-Actinins, ALP/ Enigma family of proteins including LBD3/ZASP. In another study muscle biopsies of proband with a novel *ZASP* mutation (c.76C>T; p.Pro26Ser) revealed ZASP-positive protein aggregates co-stained with ubiquitin, p62, and LC3, along with autophagic vacuoles, indicating impaired protein turnover and activation of the autophagy pathway suggesting that this PDZ-domain mutation causes sarcomeric protein misfolding, leading to aggregate accumulation and altered autophagic response, thereby contributing to the pathogenesis of myofibrillar myopathy (Cassandrini et al. 2021).

The C-terminal end of LDB3 protein harbors three LIM domains (LIM1; aa 441-492, LIM2; aa 500-551, LIM3; aa 551-612). The LIM domains contain conserved Zinc finger motif with cystine-rich consensus sequences. Like PDZ domain, LIM domains are also protein-protein interaction domain and perform diverse biological role including regulation of actin structure and dynamics, integrin dependent cell adhesion and signaling and tissue specific gene expression. The C-terminal LIM domain of LBD3/ZASP has been shown to interact with PKC (Arimura et al., 2004). In the present study, we have detected 3 variations in the C-terminal region harboring LIM domains. The variant p.F465I is located in first LIM domain, while another (p.F496L) is present in the near vicinity of second LIM domain and the last one (p.P606S) in the third LIM domain. *In vitro* studies using C2C12 myoblast have demonstrated significant reduction in mutant protein expression compared to wild-type. Immunostaining has also revealed disruption of actin cytoskeleton with relatively weak expression of mutant protein. The downregulation of *LDB3* was also further confirmed in three of these variants at mRNA level (data not shown). Remarkable difference in predicted secondary and tertiary structure of mutant proteins compared to wild-type, showing high RMSD values for the variant p.Q402P (0.941), p.F496L (0.766) and p.P606S (0.916), suggests structural and functional impairment of mutant proteins. Down-regulation of Cypher has been reported to be associated with DCM phenotype in Cypher knockdown rat (Xuan *et al*. 2020). These authors (2020) have shown that the down-regulation of Cypher induced apoptosis in cardiomyocytes via inhibiting Akt dependent pathway and enhancing p38 MAPK phosphorylation.

Apart from this, we also identified seven synonymous variants which were already reported in dbSNP database, however we didn’t come across any published literature associated with these variants. We adopted a comprehensive *in silico* approach and identified that synonymous variants affected the conformation of mRNA structure analyzed for both 50 and 151 nucleotide structure, stability of the predicted structures and codon usage. Our Mfold based analysis suggested that note-worthy changes in the conformation rearrangement of nucleotides in case of variants c.273G>A, c.504T>C, c.546T>C, c.1041C>A and c.1074C>T. However, in case of variants c.456G>A and c.837GCC>GCT we observed no differences in the conformation of mRNA. We also examined the stability of predicted mutant mRNA structure which was decreased in case of variants c.273G>A, c.504T>C, c.546T>C, c.837C>T irrespective of nucleotide length. However, in case of variant c.1041C>A (p.S347S) the stability of predicted mRNA increased irrespective of nucleotide length. Similar analysis was performed for variant c.456G>A and c.1074C>T which revealed that the stability for 151 nucleotide structure decreased and increased respectively. Since mRNAs regulate protein expression any changes in the structure or stability of mRNA can alter protein expression. The protein synthesized per mRNA molecule is also a function of how well the translation machinery initiates and elongates on the coding sequence which in turn is dependent on mRNA structure and stability (Mauger *et al*. 2019). The characterization of transcriptome-wide RNA structure has also revealed a correlation between the structure of mRNA and levels of protein expression (Ding *et al*. 2014). It is not only the mRNA characteristics which affects the protein expression, translational kinetics, and co-translational protein folding but these characteristics are also governed by differences in the codon usage frequencies. It is also noteworthy that differential usage of synonymous variants due to variable speed of decoding on ribosome may be suggestive of a “code” within the genetic code for operating accurate co-translational protein folding and subsequently, maintenance of cellular functions (Koutmou *et al*. 2015; Mitra *et al*. 2016). We also analyzed the codon usage of synonymous variants adopting the relative synonymous codon usage method and codon usage which indicated differences in the preference of codon across wild-type and mutant. These two approaches suggested that in case of variants c.273G>A (p.T91T), c.456G>A (p.A152A), c.504T>C (p.D168D), c.546T>C (p.S182S), mutant codon is more frequently used than wild-type codon whereas in case of variants c.837C>T (p.A279A), c.1041C>A (p.S347S) and c.1074C>T (p.A358A) wild-type codon is more commonly used when compared to mutant codon.

## Conclusion

Thus, taken altogether our study supports the role of *LDB3* in maintaining the structure and function of Z-disc, which is evident by our protein expression and cellular localization studies. The high mutation frequency of 21% also suggests its possible association with the DCM phenotype which also enables it to be used as a biomarker in Indian population. The identified mutations also exhibited phenotypic heterogeneity. The characterization of some of the selected variants that have shown aberrant expression at protein level have been characterized using global genome-wide and proteome-wide approaches (unpublished data) revealing differential expression of many of calcium ion, cardiac muscle contraction and development related genes and proteins.

## Acknowledgment

We deeply appreciate the patients and their families for their invaluable contributions to this study. We are also profoundly thankful to Padma Shri Prof. T.K. Lahiri and Prof. D. Agrawal from the Department of Cardiothoracic & Vascular Surgery, IMS, BHU, Varanasi for their constant encouragement and valuable support in enrolling patients for the study.

