## Supplementary data for "Identification and functional characterization of *LDB3* gene variants in DCM patients from Indian Population"

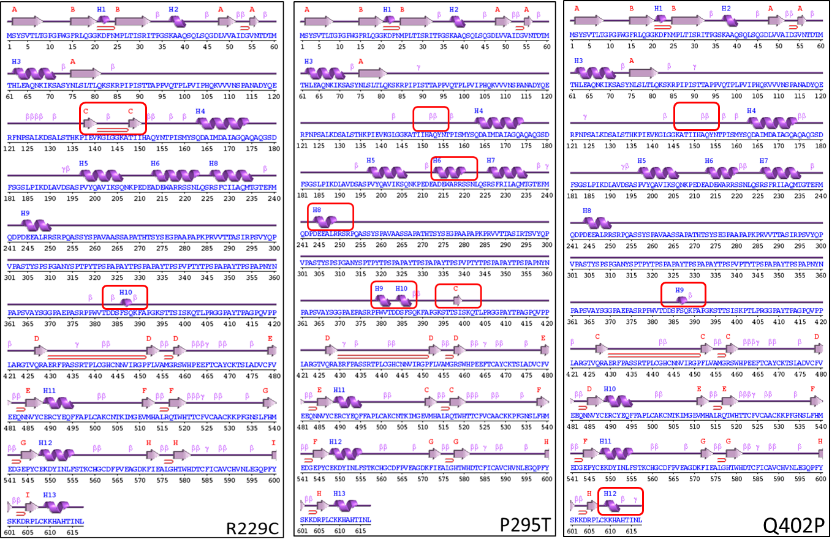

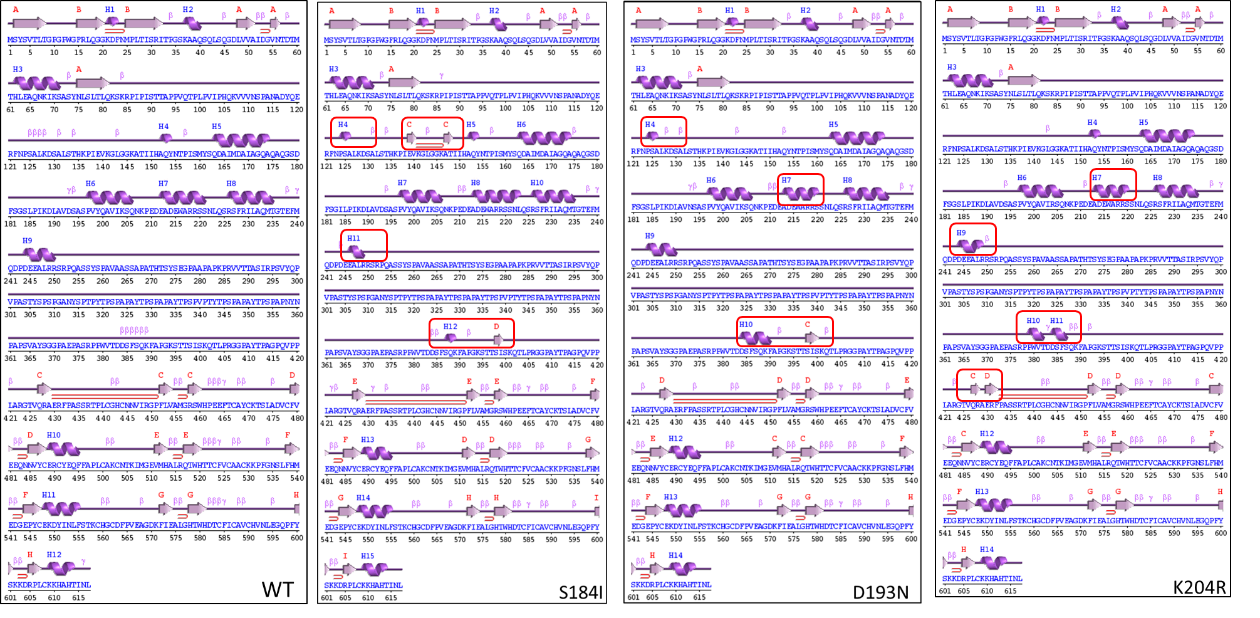

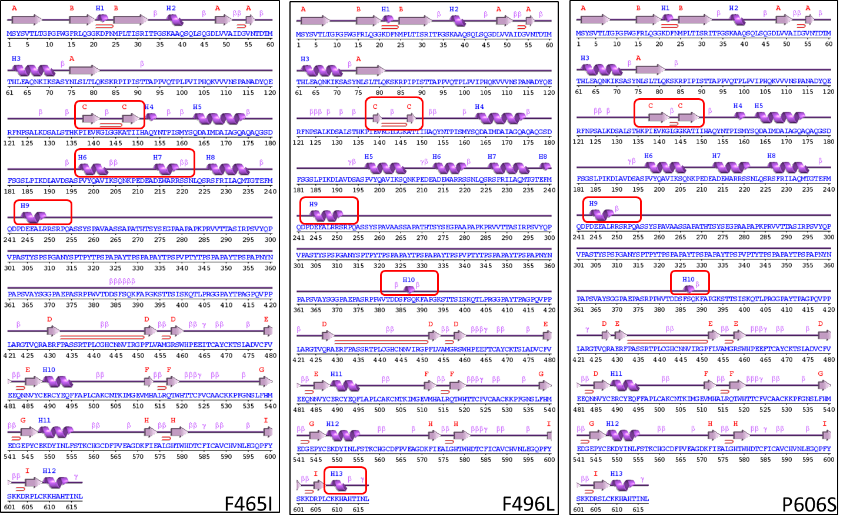

***Supplementary Figure 1.*** *Secondary structure of LDB3 protein predicted for wild-type (WT) and the variants by PDBsum tool*

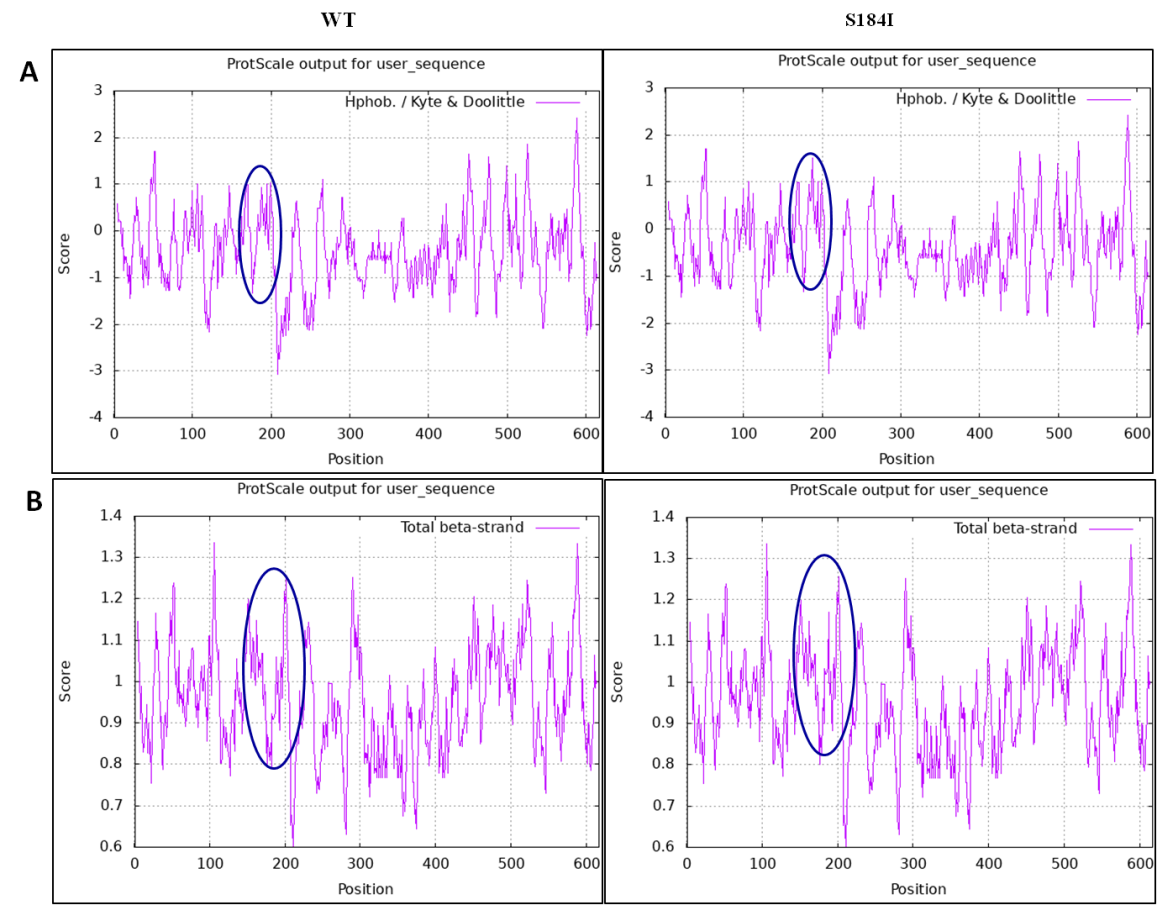

***Supplementary Figure 2:*** *The profile for physicochemical properties of variant p.S184I. (A) Hydrophobicity (B) Total β-strand.*

**Supplementary Table 1. List of intronic variants identified in *LDB3***

| Genetic variant | No. of case (s) (MAF) | No. of control (s) | INDEX-db | IndiGen*omes* | rsID/RCV  (MAF) |
| --- | --- | --- | --- | --- | --- |
| c.322-50G>C | 2/100 (0.01) | 0/100 | NR | 0.7895 | rs3740345 (0.375) |
| c.548+53A>C | 1/100 (0.005) | 0/100 | NR | 0.0273 | rs111941601 (0.043) |
| c.718+47G>C | 13/100 (0.065) | 0/100 | NR | 0.2134 | rs3740346 (0.199) |
| c.718+81G>A | 8/100 (0.04) | 0/100 | NR | NR | Novel  (RCV001293348.1) |
| c.755+11G>A | 1/100 (0.005) | 0/100 | NR | NR | Novel  (RCV001293349.1) |
| c.755+86G>A | 1/100 (0.005) | 0/100 | NR | NR | Novel  (RCV001293350.1) |
| c.*30C>G | 1/100 (0.005) | 0/100 | NR | NR | Novel  (RCV001293351.1) |

**Supplementary Table 2.** Changes in the secondary structure of the LDB3 protein due to the non-synonymous variants identified in the present study

| Changes In Secondary Structure | | Variants | | | | | | | | |
| --- | --- | --- | --- | --- | --- | --- | --- | --- | --- | --- |
|  |  | **p.S184I** | **p.D193N** | **p.K204R** | **p.R229C** | **p.P295T** | **p.Q402P** | **p.F465I** | **p.F496L** | **p.P606S** |
| Introduction of Α-Helix | **Amino Acid Residues** | 124-125  387-389 | 124-125  383-390 | 379-382  384-387 | 387-389 | 379-381  384-387 | 387-389 | - | 387-389 | 387-389 |
| Loss of Α-Helix |  | - | - | - | - | 152-154 | 152-154 | - | - | - |
| Lengthening of Α-Helix |  | - | - | - | - | - | - | - | 250-251 | - |
| Reduction of Α-Helix |  | 248-250 | 219-221 | 219-221  250-251 | - | 219-221  249-250 | 609-612 | 203-206  219-221  249-250 | 609-612 | 249-250 |
| Introduction of β-Sheet |  | 139-140  147-148  398-399 | 398-401 | 430-432 | 139-140  147-148  398-399 | 397-399 | - | 138-141  147-150 | 139-140  147-148  398-399 | 138-142  145-149 |
| Introduction of β-hairpin |  | 141-146 | - | - | 141-146 | - | - | 142-146 | 141-146 | 143-144 |
| Loss of β-Sheet |  | - | - | - | - | - | - | - | - | - |
| Lengthening of β-Sheet |  | - | - | - | - | - | - | - | - | - |
| Reduction of β-Sheet |  | - | - | - | - | - | - | - | - | - |

**Supplementary Table 3**: List of so far published mutations, identified in ZASP

| **Variant** | **Cardiomyopathy Phenotype** | **MAF on GnomAD** | **Location** | **References** |
| --- | --- | --- | --- | --- |
| p.Gly20Alafster41 | DCM | 1.860 x10^-6^ | PDZ motif | Koopmann *et al*. 2023 |
| p.Pro26Ser | MFM | Absent | PDZ motif | Cassandrini *et al*. 2021 |
| p.Asp117Asn | DCM, LVNC | 7.10 x 10^-3^ | - | Vatta *et al.* 2003 |
| p.Lys136Met | DCM, LVNC | Absent | - | Vatta *et al.* 2003 |
| p.Ala147Thr | MFM | Absent | - | Selcen *et al.* 2005 |
| p.Asn155His | Distal MFM | Absent | DUF domain | Zheng *et al*.2016 |
| p.Ala165Val | DCM | Absent | DUF4749 | Martinelli *et al.* 2014 |
| p.Ala171 Thr | DCM | 1.51 x 10^-4^ | DUF4749 | Martinelli *et al.* 2014 |
| p.Ser189Leu | DCM | 5.19 x 10^-4^ | DUF4749 | Vatta *et al.* 2003 |
| p.Ser196Leu | DCM, LVNC, HCM | 6.57 x 10^-6^ | DUF4749 | Vatta *et al.* 2003 |
| p.Lys204Arg | DCM, LVNC | 1.56 x 10^-2^ | DUF4749 | Vatta *et al.* 2003 |
| p.Thr206Ile | DCM | 6.57 x 10^-6^ | DUF4749 | Vatta *et al.* 2003 |
| p.Thr213Ile | DCM, LVNC | Absent | DUF4749 | Vatta *et al.* 2003 |
| p.Ala222Thr | MFM | 3.48 × 10^−4^ | DUF domain | Weihl *et al*. 2015 |
| p.Asp261Asn | Distal Myopathy | 6.195× 10^−7^ | - | [Gadaleta](https://pubmed.ncbi.nlm.nih.gov/?term=%22Gadaleta%20G%22%5BAuthor%5D) *et al*. 2025 |
| p.Arg268Cys | MFM | 4.61 × 10^−5^ | - | Selcen *et al*. 2005 |
| p.Ile345Met | DCM | Absent | Atrophin-1 | Vatta *et al.* 2003 |
| p.Ile352Met | DCM | Absent | Atrophin-1 | Vatta *et al.* 2003 |
| p.Asp366Asn | HCM | Absent | Atrophin-1 | Theis *et al.* 2006 |
| p.Val407ThrfsTer84 | DCM | 6.200× 10^−7^ | Atrophin1 | Koopmann *et al*. 2023 |
| p.Gln414Lys | MFM | 4.60 × 10^−5^ | Atrophin1 | Weihl *et al*. 2015 |
| p.Tyr468Ser | HCM | Absent | LIM 1 domain | Theis *et al.* 2006 |
| p.Gln519Pro | HCM | Absent | LIM 2 domain | Theis *et al.* 2006 |
| p.Arg541* | DCM | 6.198× 10^−6^ | LIM 2 domain | Koopmann *et al*. 2023 |
| p.Val566Met | Distal dominant weakness | 2.63 × 10^−5^ | LIM 3 domain | Cai *et al.* 2012 |
| p.Pro615Leu | HCM | Absent | LIM 3 domain | Theis *et al.* 2006 |
| p.Asp626Asn | DCM | Absent | LIM 3 domain | Arimura *et al.* 2004 |
| p.Gln627Hisfster75 | DCM | Absent | LIM 3  domain | Koopmann *et al*. 2023 |
